# Cathelicidin downregulates neurotensin and substance P hippocampal levels

**DOI:** 10.1101/2023.07.07.548146

**Authors:** Ismael Perez Flores, Suely Kubo Ariga, Hermes Vieira Barbeiro, Denise Frediani Barbeiro, Fabiano Pinheiro da Silva

## Abstract

**Background:** Sepsis is a life-threatening condition and septic encephalopathy is an early and frequent manifestation of this disease. Antimicrobial peptides are important components of innate immunity playing a crucial role during bacterial infections. Here, we investigate the protein levels of several neuropeptides in CRAMP-deficient and wild-type mice, in healthy conditions and following experimental sepsis.

**Methods:** Mice were submitted to cecal ligation and puncture and the protein levels of neurotensin, substance P, oxytocin and β-endorphin were evaluated in the brain.

**Results:** We found that CRAMP-deficient mice produce significantly less neurotensin and substance P than wild-type mice in the hippocampus, both before and 24 hours following experimental sepsis, but not 15 days post-septic shock.

**Conclusions:** The hippocampus is a complex structure, highly vulnerable during sepsis. The role of antimicrobial peptides and their interplay with neuropeptides should be further evaluated in this scenario.

## 1. Introduction

Sepsis is a life-threatening disease attributed to a deregulated host response to infection. About half of sepsis survivors, moreover, show a significant decrease in quality of life and a substantial increase in mortality after hospital discharge, when compared with the overall population [1].

Cognitive dysfunction is one of the most common chronic complications of septic shock [2]. Symptoms include neurocognitive impairment and psychiatric disorders that can persist for years. Markers of neuroinflammation, such as NSE, Tau protein, β-Amyloid protein and S100β protein have been investigated to diagnose sepsis encephalopathy and to monitor potential therapies. Using experimental models, high levels of these proteins have been found in the brain tissue, showing a positive correlation with inflammatory cytokines [3].

Neuropeptides participate in complex and different aspects of physiology, including sexual behavior, mood, circadian rhythm, regulation of food intake and inflammatory balance [4].In a prospective study, decreased plasma levels of neurotensin, substance P, oxytocin and α-MSH levels were detected in the presence of septic shock [5].

Cathelicidins are endogenous substances that play an important role in innate immunity [6]. Humans and rodents produce only one cathelicidin family member named, respectively, LL-37 and CRAMP. A study from our group demonstrated decreased LL-37 gene expression during septic shock, in comparison with non-infected critically ill patients or with critical patients already recovering from sepsis [7].

In the same direction, we showed that CRAMP-deficient animals exhibited lower mortality and increased levels of several inflammatory cytokines, when submitted to experimental sepsis, putting in evidence that lower levels of cathelicidins may be beneficial in this situation [8], even though detrimental in other inflammatory diseases. Indeed, cathelicidins have been shown an intriguing dual role in systemic inflammation [9]. For these reasons, we decided to investigate the late inflammatory effects of septic shock in the brain of CRAMP-deficient and wild-type mice, by measuring the protein levels of several neuropeptides.

## 2. Material and methods

Neurotensin, β-endorphin, oxytocin and substance P protein levels were measured in the prefrontal cortex and in the hippocampus of CRAMP-deficient and wild-type animals, 24 hours or 15 days after experimental sepsis (cecal ligation and puncture). A third group of CRAMP-deficient and wild-type mice were used as healthy controls. Results were analyzed using the Kruskal–Wallis test followed by the Mann–Whitney U test. A p-value ≤0.05 was considered significant.

## 3. Results and Discussion

We found that CRAMP-deficient mice produce significantly less neurotensin and substance P than wild-type mice in the hippocampus, both before and 24 hours following experimental sepsis, but not 15 days post-septic shock (Figure 1A and 1B). No difference was detected in regard with the protein levels of oxytocin or β-endorphin in the hippocampus (Figures 1C and 1D) and no difference was detected in the prefrontal cortex among the study groups, when the protein levels of the same four neuropeptides were evaluated (data not shown).

**Figure 1.**
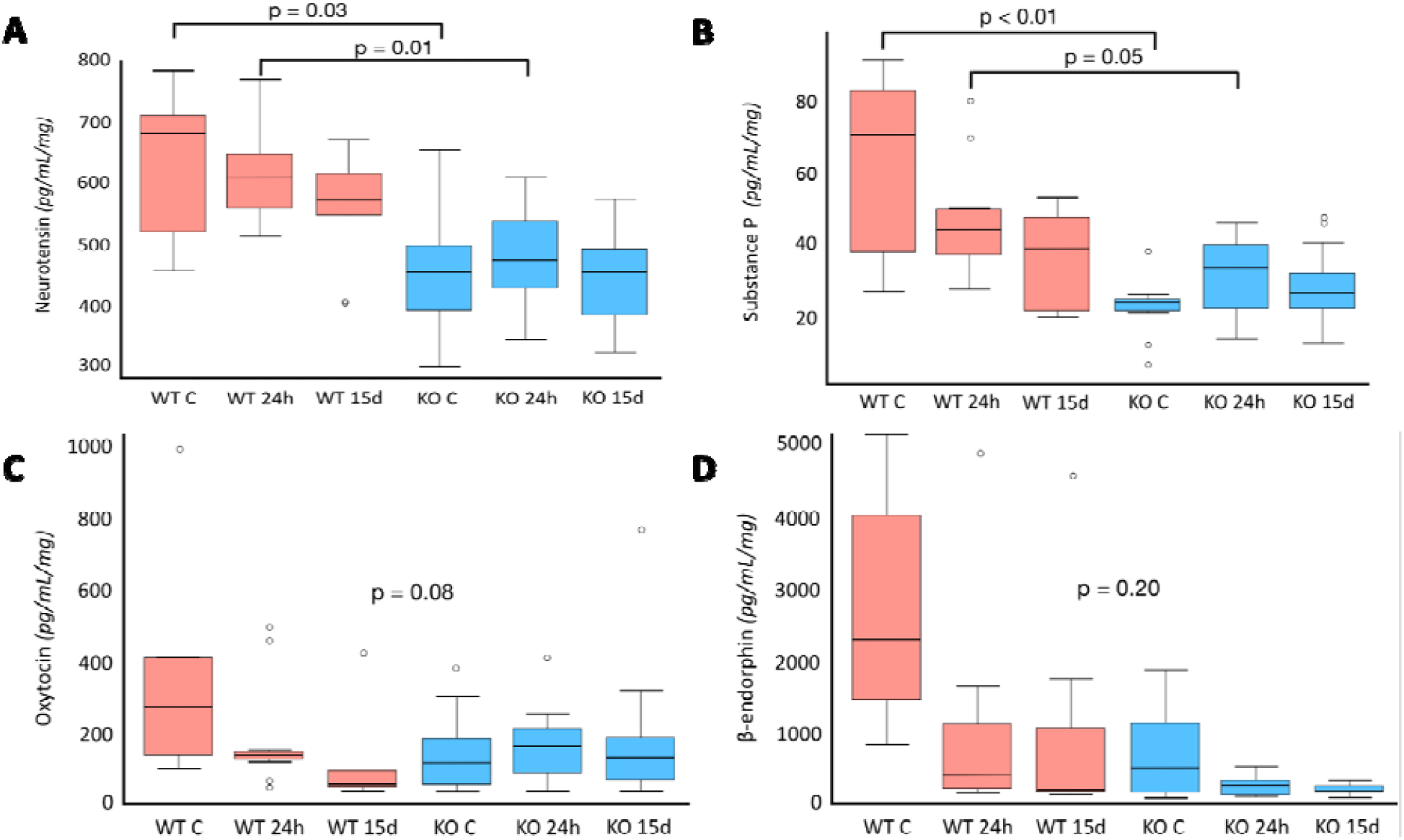
Protein levels of neurotensin, substance P, β-endorphin and oxytocin in the hippocampus of CRAMP-deficient and wild-type mice. WT C and KO C = wild-type and CRAMP-deficient healthy mice, WT 24h and KO 24h = wild-type and CRAMP-deficient mice 24 hours following experimental sepsis, WT 15d and KO 15d = wild-type and CRAMP-deficient mice 15 days after experimental sepsis (9 to 14 animals per group).

Neurotensin and substance P appear to be detrimental in sepsis [10, 11]. Among its multiple effects, neurotensin induces hypotension. Substance P plays an important role in nociception, immunity and inflammation [12]. Our results corroborate our previous publications on this subject, expanding the effects of cathelicidins in systemic inflammation. It is intriguing, however that the impact of CRAMP could be detected only in the hippocampus.

## 4. Conclusion

The hippocampus is a complex, plastic and vulnerable brain structure that plays a major role in learning and memory. The hippocampus appears to be especially vulnerable in sepsis, since survivors develop pronounced hippocampal atrophy [13]. Systemic inflammation can result in impairment of memory that persists long after the resolution of inflammation. The role of cathelicidins, neurotensin and substance P should be further investigated in this scenario.

## Availability of Data and Materials

All data and technical details are available upon request.

## Author Contributions

FPS designed the research study. IPF, SKA, DFB and HVB performed the research. FPS and IPF analyzed the data. FPS wrote the manuscript. All authors contributed to editorial changes in the manuscript.

## Ethics Approval and Consent to Participate

Protocols were in accordance with the University of São Paulo Faculty of Medicine Ethical Committee (project number 1179/2018). All authors read and approved the final manuscript.

## Acknowledgments

None.

